# Micro-/nano-plastics accentuate Parkinson’s Disease-relevant phenotypes in a *Drosophila* model

**DOI:** 10.1101/2025.11.17.688805

**Authors:** Simon A. Lowe, James E.C. Jepson

## Abstract

Micro- and nano-plastic (MNP) particles are a ubiquitous environmental contaminant that are increasingly bioaccumulating in human tissues, particularly the brain. MNPs induce mitochondrial defects, oxidative stress, inflammatory responses and neurotoxicity in cellular and organismal models. This raises the possibility that MNP exposure could cause or exacerbate neurological conditions associated with these pathological phenomena. Parkinson’s Disease (PD), a common movement disorder characterised by degeneration of striatal dopaminergic neurons, and associated with mitochondrial dysfunction, represents such a condition. We therefore hypothesised that MNP exposure might interact with PD risk mutations affecting mitochondrial fidelity. We used a fruit fly model of *PRKN*-dependent PD associated with defects in mitophagy, a mitochondrial quality control pathway, to test this hypothesis. We found that ingestion of MNPs at concentrations tolerated by wild-type controls selectively enhanced PD-relevant phenotypes – including progressive dopaminergic neurodegeneration, movement defects, and sleep disruption – in this model of PD. Our data suggest that defects in mitochondrial quality control can increase vulnerability to MNP exposure, and more broadly, that MNPs may synergistically interact with existing genetic risk factors to worsen neurological disease.

## INTRODUCTION

The rapidly changing environment has the potential to impact human neuronal health via increasing global temperatures ^1^, reduced air quality ^2^, and exposure to neurotoxic environmental pollutants ^3^. Collectively, neurological disorders are currently the leading cause of disability worldwide; and incidence rates and associated social challenges are expected to increase in the coming decades ^4^. Due to the ageing global population, this increase is projected to be particularly prominent for age-related neurological disorders such as Parkinson’s Disease (PD) ^5^. Thus, understanding how the changing environment may impact the incidence and progression of these disorders is essential to inform policy and shape social responses.

The impact of plastic pollution on human health ^6^, and particularly neurological health ^7^ ^8^, is emerging as a potential global health crisis. Micro- and nano-plastics (MNPs) are defined as plastic particles on the ≤ 1 μM and 1 μM - 1 mm scale respectively, which are either manufactured or generated from the breakdown of macro-plastics by UV light degradation, mechanical abrasion from water and sand, etc ^9^. Global annual plastic production exceeds 400 million tons, a majority of which are single-use, and only a small minority of which is recycled ^10^. MNP pollution is now ubiquitous: significant levels have been identified in the air, soil, ocean and drinking water, including in remote regions ^9^, with evidence that they bioaccumulate up the food chain ^11^. Unsurprisingly given the range of absorption routes, MNPs have been isolated in a range of human tissues ^12^. MNPs cross the blood-brain barrier (BBB) and preferentially bioaccumulate in the brain over other tissues ^13^. Levels in post-mortem human brain tissue positively correlate with more recent time of death (2016 vs 2024) ^12^, consistent with increasing environmental levels. Particularly worryingly, MNPs have been identified in the placenta ^14^, indicating that nervous system exposure to constantly rising MNP levels from the earliest stages of development onwards will now be the norm. The health implications of this exposure are not understood, but MNP levels were higher in brains with documented dementia diagnoses ^13^, suggesting a possible causative role and indicating an urgent need to understand how MNPs interact with neural tissue and organismal health.

An extensive body of work demonstrates the cytotoxicity of MNPs to cultured cells. Exposure disrupts the morphology and function of mitochondria ^15^, possibly by binding directly to complex 1 ^16^, and induces excess production of reactive oxygen species (ROS), oxidative stress ^17^, and upregulation of inflammatory markers ^18^. These effects overlap with cellular mechanisms known to contribute to a range of neurological disorders, particularly PD, in which the mechanistic contributions of mitochondrial dysfunction, oxidative stress, and chronic inflammation are well documented ^19^. Dopaminergic neurons, which selectively degenerate in PD, are particularly vulnerable to mitochondrial toxins ^20^, and exposure to mitotoxic pesticides may contribute to the aetiology of PD ^21^ ^22^. This raises the possibility that exposure to MNPs could similarly induce PD-like symptoms. Indeed, oral ingestion or inhalation of MNPs cause PD-like motor defects and dopaminergic degeneration ^23^ ^24^ ^25^ in mice, alongside a wide range of behavioural defects in vertebrate and non-vertebrate models^26 27 28 29^.

On the basis that vulnerable populations are likely to be first impacted by adverse conditions, we hypothesised that MNP exposure might interact with intrinsic genetic risk for PD. Thus, as environmental contamination levels rise, carriers of risk mutations would be expected to be first impacted, prefiguring effects on the general population. The PINK1/Parkin-mediated mitophagy pathway represents a potential nexus for vulnerability. This pathway performs essential mitochondrial quality control by targeting damaged mitochondria for degradation. PINK1 accumulates on the outer membrane of depolarised mitochondria, recruiting the E3 ubiquitin ligase Parkin, which (poly)ubiquitinates mitochondrial outer membrane proteins, tagging the mitochondria for engulfment by the autophagosome and transport to the lysosome for degradation ^30^. A number of the strongest genetic risk factors for PD cluster in genes encoding components of this pathway, including *PINK1, PRKN* and *DJ-1* ^31^ ^32^. It is believed that impaired mitophagy results in an accumulation of defective mitochondria, leading to excessive ROS production, oxidative stress and an inflammatory response, which contribute to dopaminergic neurotoxicity ^31^. Mitophagy is upregulated in response to mitochondrial toxins such as MPP+ and rotenone, and loss-of-function *PRKN* and *PINK1* mutations causing defective mitophagy increase vulnerability to their neurotoxic effects in patient-derived dopaminergic neuronal culture and *Drosophila* models ^33^ ^34^ ^35^. Exposure to MNPs also induces PINK1/Parkin-mediated mitophagy in differentiated dopaminergic neuronal cell culture ^36^. We therefore hypothesised that genetic defects in this pathway would increase susceptibility to MNP neurotoxicity.

We tested this hypothesis using *Drosophila* as an in vivo model. Due to its relatively short lifespan and an extensive toolkit for genetic manipulation, this species represents a powerful system to interrogate age-dependent neurodegeneration, the role of the highly conserved PINK1/Parkin pathway in PD ^37^, and genetic variance in resistance to environmental toxins associated with PD ^38^ ^39^. LOF of the highly conserved *Drosophila PRKN* homologue *parkin* (*park*) causes impaired age-related mitophagy ^40^ and recapitulates key aspects of PD ^41^ ^42^. We found that lifetime exposure MNPs in the form of orally ingested polystyrene micro-spheres accelerated dopaminergic neurodegeneration and exacerbated behavioural phenotypes in *park* LOF flies, at levels tolerated by controls. We conclude that disease-associated mutations can alter susceptibility to MNP neurotoxicity; that environmental MNP pollution may impact the incidence or progression of PD; and that genetic variations in mitophagy may define a group at particular risk from MNP exposure.

## RESULTS

### Ingested MNPs enter the brain and induce a cellular stress response in *Drosophila*

We raised and maintained control flies and flies homozygous for the well-characterised *park*^1^ LOF mutation on food contaminated with 100 parts-per-million (ppm) 100 nm-diameter polystyrene nanospheres (hereafter “MNP food”, compared to “std food”). While not faithfully recapitulating the diversity of plastic polymers present in real-world conditions, several studies have reported the effects of similar feeding protocols ^43,44,45^, and found only subtle effects on viability and behaviour. Hence, this condition allowed us to test for increased vulnerability to conditions known to be broadly tolerated by flies with intact mitophagy.

Initially, we validated the effects of this feeding protocol in control flies. While extensive evidence demonstrates that MNPs cross the blood-brain-barrier in humans and mammalian models ^13^ ^46^, it has been reported that orally-ingested microspheres fail to permeate the *Drosophila* brain ^28^. In contrast, we clearly identified ingested fluorophore-tagged MNPs in the brains of adult flies (Fig. 1A and Supp. Fig. 1). The difference between our own and prior studies is likely due to the smaller size of the MNPs used herein (0.1 vs 1-5 μM ^28^), which has been demonstrated to affect efficiency of crossing cell membranes ^47^.

**Figure 1.**
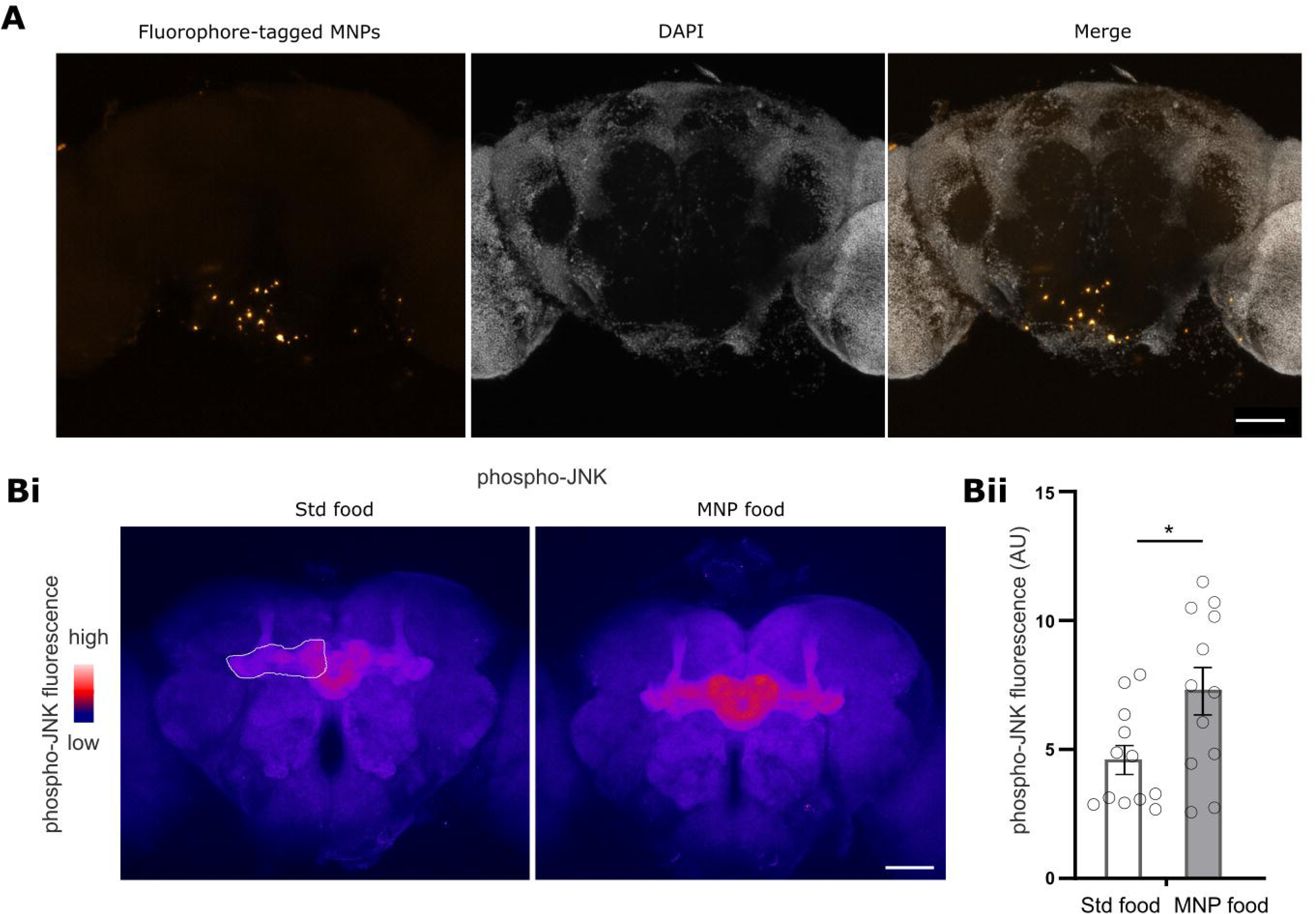
Orally ingested polystyrene microspheres permeate the brain and activate the JNK stress pathway. **A.** Representative image of a whole ex vivo brain from a 20 d post-eclosure control fly raised on food contaminated with 100ppm fluorophore (540/560)-tagged 0.1 μM polystyrene microspheres. *Left:* tagged micro-spheres; *Centre*: DAPI counterstain; *Right*: merge. Scale bar = 50 μM. **B. i.** Representative image of a whole ex vivo brain stained with anti-phospho-JNK, from a 20 d post-eclosure control fly raised on (*left*) standard food or (*right*) food contaminated with 100ppm 0.1 μM polystyrene microspheres (MNP). Representative ROI around horizontal mushroom body (MB) lobes is indicated. Scale bar = 50 μM. **ii.** Quantified anti-phospho-JNK immunofluorescence within horizontal (MB) lobes (two analysed per brain). Comparison is Mann-Whitney test, *P ≤ 0.05.

Activation via phosphorylation of the c-Jun N-terminal kinase (JNK) is a primary response to ROS production ^48^ ^49^. At 20 d post-eclosure, we observed increased levels of phospho-JNK within the horizontal mushroom body lobes - a neuropil region of the brain which receives dopaminergic input ^50^ and exhibited robust phosphor-JNK immunofluorescence - in flies raised on MNP food (Fig. 1B), confirming that our protocol replicates the extensively reported finding that MNP exposure induces excess ROS production and oxidative stress in a range of model systems ^49^ ^51^ ^52^. We did not quantify volume or spatial distribution of MNPs in the brain, but note that MNP localisation does not appear to overlap with regions of high phospho-JNK (Fig. 1B and Supp. Fig. 1), suggesting a systemic stress response.

### MNP exposure accelerates degeneration of dopaminergic neurons in *park* mutants

The ∼ 130 dopaminergic neurons in the adult *Drosophila* brain are well-characterised and readily visualised by immunostaining against the dopamine precursor tyrosine hydroxylase (TH) (Fig. 2A) ^50^. *park* LOF causes selective progressive degeneration of the protocerebral posterior lateral 1 (PPL1) subpopulation, while surrounding populations are spared ^42^. PPL1 neurons are considered to share homology with dopaminergic neurons of the mammalian substantia nigra pars compacta region ^53^, which selectively degenerate in PD and appear particularly susceptible to mitochondrial damage and oxidative stress ^54^. As expected ^42^, aged (30 d post-eclosure) but not young (3-5 d) *park*^1^ flies raised on std food exhibited an ∼ 20% reduction in TH^+^ cell bodies within the PPL1 cluster. Notably, this trend was exacerbated by lifetime exposure to MNPs (Fig. 2B,C), with no cell loss observed in two other distinct TH^+^ subpopulations (Supp. Fig. 2). These data suggest that MNP exposure does not cause widespread neurodegeneration, but instead exacerbates specific phenotypes induced by *park* LOF. No loss of neurons in any subpopulation was observed in control flies (Fig. 2B-C, Supp. Fig 2), further supporting the premise that defective Parkin-dependent mitophagy increases susceptibility to MNP neurotoxicity.

**Figure 2.**
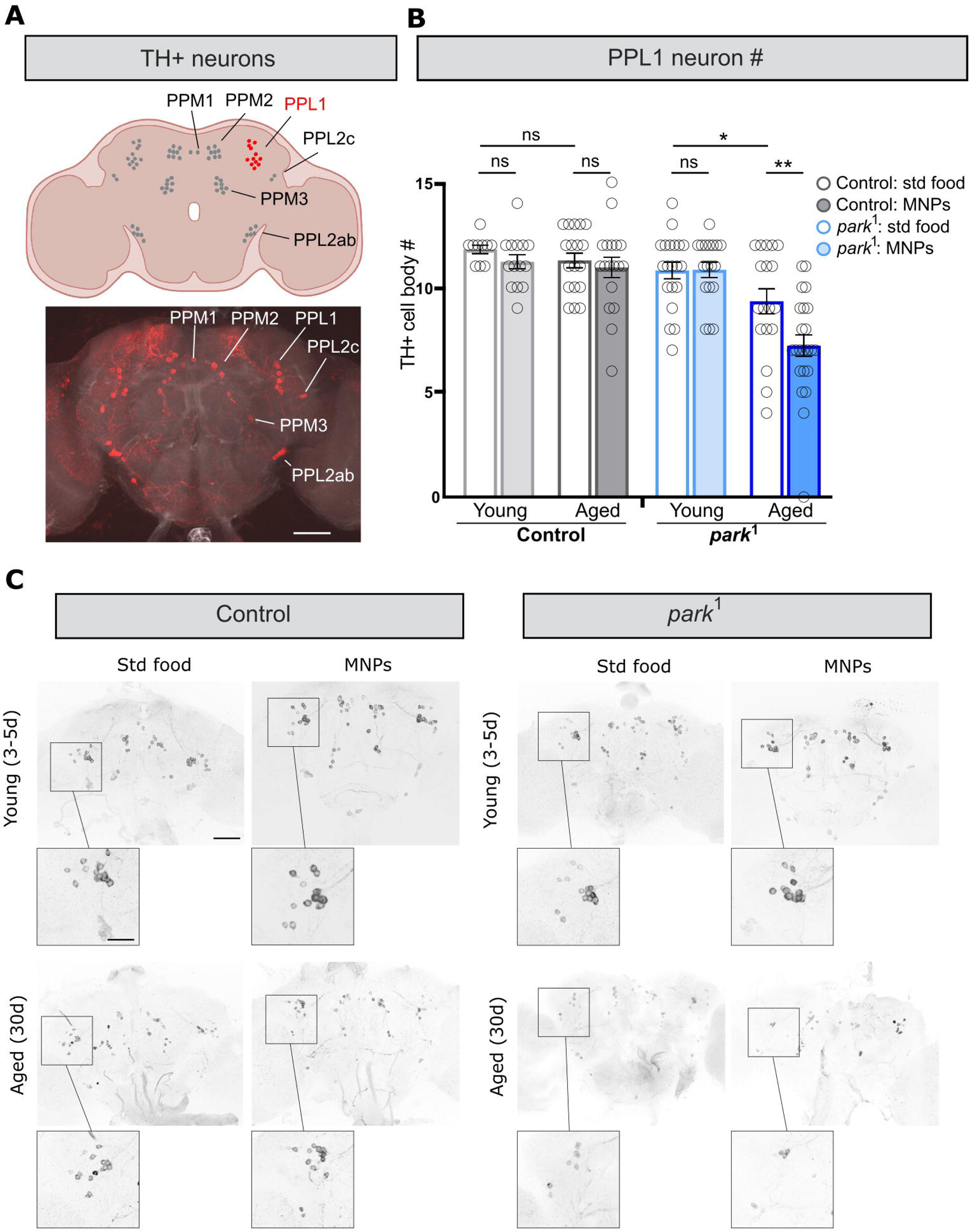
Chronic MNP exposure accelerates degeneration of dopaminergic PPL1 neurons in *park*^1^ mutants, at dosages tolerated by controls. **A.** *(Top)* Schematic showing the position of the protocerebral posterior medial (PPM) 1-3 and protocerebral posterior lateral (PPL) 1-2c clusters of TH^+^ neurons in the adult *Drosophila* brain, with the degenerating PPL1 cluster highlighted. *(Bottom)* Representative ex vivo brain (control, std food, 3 d) immunostained with anti-TH (red) against F-actin counterstain (grey). Scale bar = 50 μm. **B.** Manually counted TH^+^ cell bodies within the PPL1 subpopulation. Grey bars indicate control and blue bars *park*^1^ flies. Empty bars indicate flies raised and maintained on std food; filled bars indicate MNP-containing food. Comparison is 2-way ANOVA, within genotype, with age (young = 3-5 d; aged = 30 d) and diet (std vs MNP-containing food) as factors. *P ≤, * 0.05, **P < 0.01 **C.** Representative control and *park*^1^ ex vivo brains immunostained with anti-TH, comparing young (3-5 d) vs aged (30 d) flies, raised on std vs MNP-containing food. Enlarged insets show left PPL1 subpopulation. Scale bars = 50 μm (main), 25 μm (insets).

### MNPs exacerbate dysfunctional limb kinematics in *park* mutants

The defining clinical characteristic of PD is progressive decline in motor performance ^55^, which is recapitulated in *park* LOF mutant flies ^56^. Based on the accelerated rate of dopaminergic neurodegeneration, we hypothesised that lifetime MNP exposure would exacerbate motor decline in *park*^1^ flies. We tested gross movement in response to a mechanical stimulus at 3, 10, 20 and 30d post-eclosure, using a video-tracking method: the *Drosophila* ARousal Tracking (DART) system ^57^ ^58^. As previously reported across a range of assays ^59^ ^60^, control flies raised on standard food exhibited an age-dependent decline in motor response. Exposure to MNPs caused a transient *increase* in response in 10 d old controls, consistent with the MNP-induced hyperactivity reported in a range of model systems ^61^ ^62^ ^44^. As expected, *park*^1^ flies exhibited significantly decreased responses at all time points tested; however, MNP exposure caused no significant difference at any age (Supp. Fig 3).

We hypothesised that an MNP-induced exacerbation of movement defects might be masked by an opposing tendency towards hyperactivity, or by a ‘floor effect’, whereby aged *park*^1^ flies moved so little that enhancement of the motor phenotype was not apparent. We therefore turned to a more rigorous analysis of locomotor performance, using the Feature Learning Leg Segmentation and Tracking (FLLIT) system ^63^ ^58^ to analyse limb kinematics in high-speed videos of freely walking flies (Fig. 3A-B). First, we identified three gait parameters by which *park*^1^ flies differed from controls. These are: reduced stride length (displacement), reminiscent of the shuffling gait observed in PD ^64^; increased non-linearity of limb movement (derived by normalizing the path moved by a limb during a stride to the distance displaced), indicative of a loss of motor co-ordination; and reduced % of time a limb was moving, indicative of a “plodding gait” with greater pauses between limb movements.

**Figure 3.**
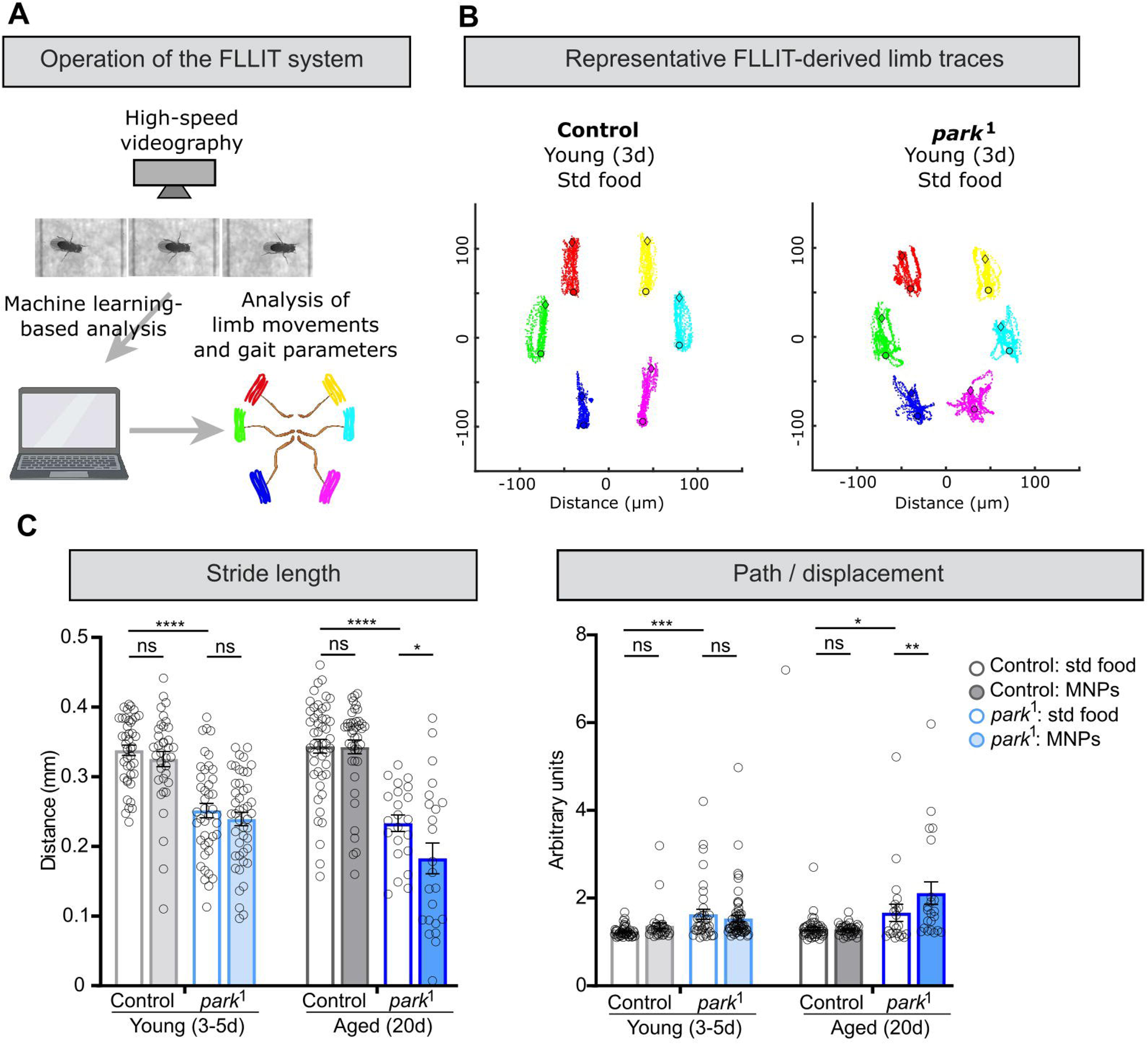
Chronic MNP exposure exacerbates dysfunctional limb kinematics in *park*^1^ mutants, at dosages tolerated by controls. **A.** Schematic showing the function of the Feature Learning-based Limb segmentation and Tracking (FLLIT) system for analysing limb kinematics. **B.** Representative FLLIT-derived schematics showing overlaid movement of each limb over multiple strides, relative to body center, for control (left) and *park*^1^ 3 d old flies. **C.** Quantification of two FLLIT-derived gait parameters: stride length (displacement) and path/displacement. Data points represent individual limbs, 6 per fly. Comparisons are 2-way ANOVA, within age point, with genotype and diet as factors. *P < 0.05, **P < 0.01, *** P < 0.001, ****P < 0.0001.

Testing at 20 d (since by 30 d *park*^1^ flies moved very little), we found that long-term MNP exposure exacerbated the reduced displacement and enhanced non-linearity of movement in *park*^1^ flies, with no effect on controls (Fig. 3C). The effect on movement % was more complex. Young *park*^1^ exposed to MNPs exhibited an increased movement % relative to those raised on standard food, with controls showing a similar, albeit non-significant, trend (p = 0.08). By 20 d, however, MNP-exposed control flies exhibited increased movement % relative to those raised on standard food, while the effect had disappeared in *park*^1^ flies (Supp. Fig. 4).

We conclude that MNP exposure indeed causes hyperactivity, manifesting here as more rapid limb movements while walking, while simultaneously exacerbating the *park*^1^ phenotype of a slower, plodding gait. Thus, *park*^1^ flies are affected in a qualitatively different manner to controls; first, hyperactivity becomes apparent after shorter periods of exposure due to their enhanced susceptibility to the neurotoxic effects of MNPs, then by the time hyperactivity manifests in controls, the effect is masked by exacerbation of the opposing phenotype of a reduction in movement %.

### MNP exposure exacerbates night sleep loss in *park*^1^ mutants

Sleep dysfunction is increasingly recognised as a feature of PD, both as a primary symptom affecting quality of life, and as a predictor of and potential factor contributing to disease progression ^65^. In *Drosophila*, *park* LOF causes circadian abnormalities and sleep disturbance ^66^ ^67^. We interrogated the effect of MNP exposure on this phenotype using 5H *Drosophila A*ctivity *M*onitors (DAM5H) (Fig. 4A), a multi-infrared beam system that facilitates high-resolution measures of sleep and activity in *Drosophila* ^68^. While we clearly saw an effect of *park*^1^ on both day and night sleep (Fig. 4A), we focused on the previously published phenotype of reduced night sleep ^66^. Strikingly, in *park*^1^ flies, MNP exposure caused a further reduction in night sleep levels by 20 d, with no significant effect observed in control flies (Fig. 4A, B).

**Figure 4.**
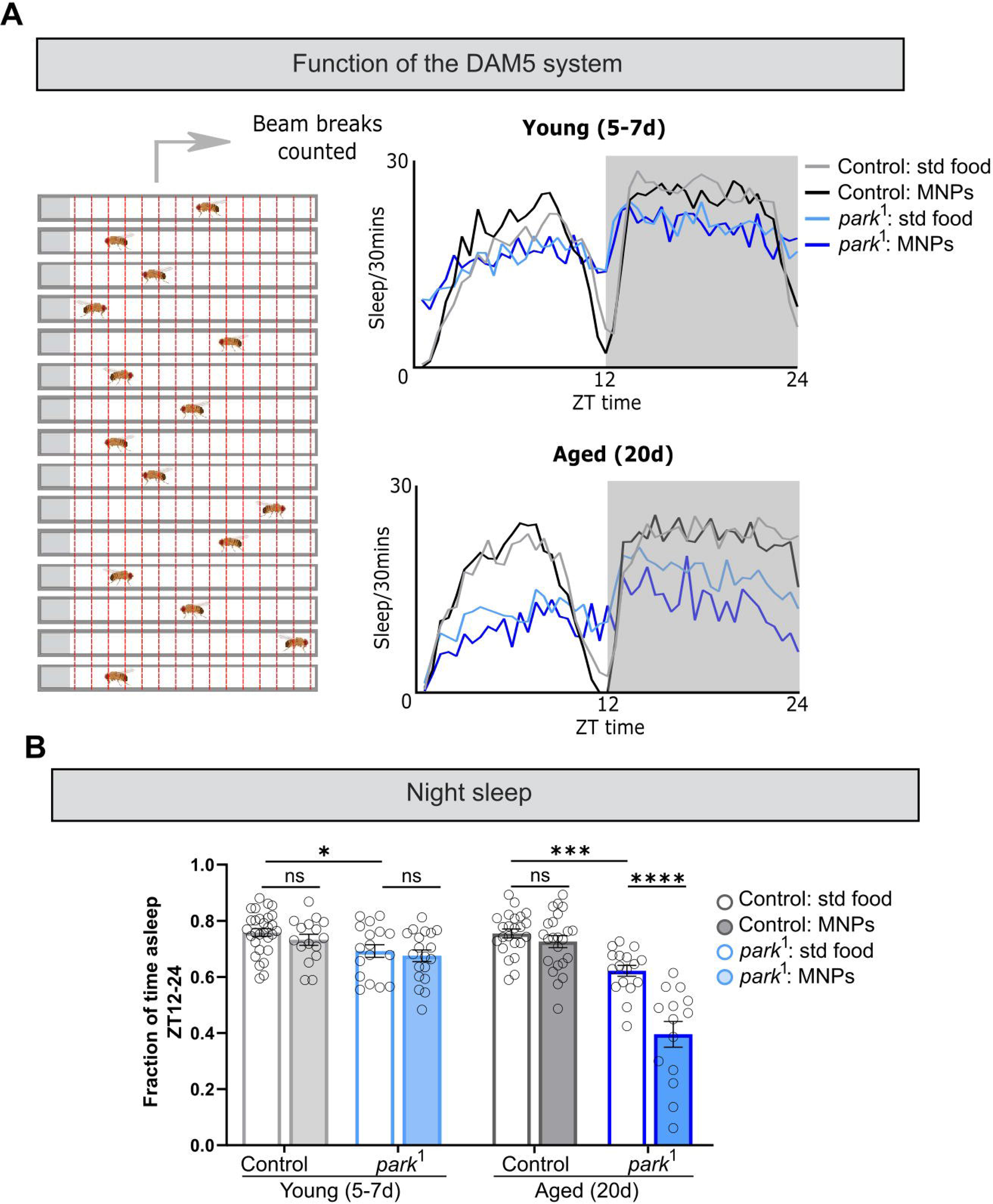
Chronic MNP exposure exacerbates night sleep loss in *park*^1^ mutants, at dosages tolerated by controls. **A.** *(Left)* Schematic showing function of the DAM5 system. *(Right)* Mean sleep in 12L: 12D conditions of young (5-7 d) and aged (20 d) *park*^1^ and control flies, raised on standard or MNP food. **B.** Night sleep, as the fraction of time from ZT12-24 (lights-off) spent asleep. Data points represent individual flies. Comparisons are 2-way ANOVA, within age point, with genotype and diet as factors. *P < 0.05, *** P < 0.001, ****P < 0.0001.

Collectively, the above data demonstrate a detrimental and genotype-specific impact of chronic MNP on disease-relevant phenotypes in a well-characterized *Drosophila* model of PD.

## DISCUSSION

MNP pollution is now ubiquitous in the environment, and bioaccumulation in the human brain, together with a wealth of evidence of cytotoxic effects from *in vitro* and *in vivo* models, raise the possibility that chronic exposure could cause or contribute to neurological disease ^6 7^. Understanding how this environmental factor interacts with underlying genetic vulnerabilities can therefore help delineate the scale of the problem and identify populations at specific risk.

Together, the data presented herein demonstrate that chronic MNP exposure exacerbates degenerative, motor, and non-motor PD-like phenotypes in an established invertebrate model of *PRKN*-linked PD. We conclude that disease-associated mutations can modify susceptibility to the neurotoxic effects of MNP exposure, that genetic defects in mitophagy may define a subpopulation at particular risk, and that research into MNP pollution as an environmental factor potentially impacting rising rates of PD ^5^ is warranted.

Here, we focused on PD, on the basis that the potential of environmental factors to contribute to PD-like phenotypes is well-established ^3^ ^69^; that mitotoxic effects of MNP exposure ^15^ overlap with the mechanism of action of pesticides associated with PD-like phenotypes ^22^ ^70^; and that chronic exposure has been reported to induce dopaminergic neurodegeneration in model systems ^23^ ^24^ ^25^. However, the effects of MNP exposure – such as mitochondrial dysfunction, oxidative stress and neuroinflammation ^15^ – overlap with a cellular mechanisms contributing to a range of neurodegenerative and neurodevelopmental disorders ^71^ ^72^ ^73^. Thus, the contribution of plastic pollution to a much wider range of neurological health outcomes should be considered.

Here we focused on PD-associated defects in PINK1/Parkin-dependent mitophagy as a genetic risk factor. However, the cellular effects of MNP exposure are widespread, and a large number of mutations affecting a range of cellular processes increase risk for PD ^74^, so genetic defects in mitophagy are likely not to be unique in increasing susceptibility to MNP exposure is unclear. Further, our interpretation that *park*^1^ flies’ increased vulnerability to MNPs is due to defective mitophagy is based on extensive published data on the effects of MNP exposure ^15^, Parkin LOF ^40^, and the interaction between Parkin LOF and exposure to mitochondrial toxicants ^35^ ^75^, but is not explicitly demonstrated here. Further work could directly interrogate the molecular mechanisms underlying the interaction between Parkin LOF and MNP toxicity by directly measuring rates of mitophagy, ROS production and accumulation of abnormal mitochondria, and/or strengthen the genetic evidence associating mitophagy with resistance to MNP neurotoxicity by testing whether LOF of other components of the PINK1/Parkin-dependent mitophagy pathway similarly increased vulnerability to MNP exposure ^37^ ^32^. Furthermore, while we demonstrate that MNPs permeated the brain, we did not interrogate the extent to which neuronal effects were attributable to MNPs within the brain versus elsewhere in the body, for example via the gut-brain axis ^76^ ^45^. These limitations can be addressed through further studies that build upon the data presented in this work.

MNPs levels are difficult to assess, due to challenges in direct measurement ^77^ huge range of particle size and type, geographical differences, and changes over time ^78^ ^79^ ^80^, leading to a large range of estimates in levels in relevant sources of absorption ^81^. The concentrations used in this study are likely to be significantly higher than those most of the global population are currently exposed to. In other ways, however, this feeding protocol may underestimate toxicity; it does not recapitulate the full range of MNP absorption routes ^82^, particle sizes ^47^ and shapes ^83^, or potentially additive interactions with other environmental contaminants ^84^ present in real-world conditions. Further, since MNPs bioaccumulate in brain tissue over time ^13^, short-lived organisms such as *Drosophila* may be expected to tolerate higher levels of pollutions than humans. Thus, while we demonstrate a clear interaction between a PD genetic risk factor and exposure to MNPs, further work is required to interrogate the effects in more ecologically relevant conditions.

## MATERIALS AND METHODS

### Experimental models and MNP exposure conditions

Experimental models were fruit flies of the species *Drosophila melanogaster*. Flies were maintained on standard fly food under 12 h: 12 h light-dark cycles (12L: 12D) at a constant 25°C. Experiments were conducted at the age indicated, on male flies. Genotypes used were *park*^1^ (Bloomington *Drosophila* Stock Centre stock number 34747) and the isogenised w^1118^ control line iso^31^ (a kind gift from Prof. Kyunghee Koh, Thomas Jefferson University). Flies were raised on a standard food medium. MNP food was generated by mixing 0.1 μM-diameter Opti-Bind polystyrene microparticles (ThermoFisher) to a concentration of 100pp into melted fly food. For visualisation, fluorophore-tagged 0.1 μM-diameter polystyrene FluoSpheres (excitation/emission 540/560) (ThermoFisher) were substituted.

### Immuno-histochemistry

Brains were dissected in phosphate-buffered saline (PBS) (Sigma Aldrich) at the time point indicated and immuno-stained as described previously ^85^. Briefly, brains were fixed by 20 min room temperature incubation in 4% paraformaldehyde (MP biomedicals) and blocked for 1 h in 5% normal goat serum in PBS + 0.3% Triton-X (Sigma-Aldrich) (PBT). Brains were incubated overnight in primary antibody at 4°C, washed three times in PBT and incubated overnight in secondary antibody at 4°C, then washed a final time before mounting in SlowFade Gold anti-fade mountant (ThermoFisher). Primary antibodies were rabbit anti-TH/tyrosine hydroxylase (Abcam, ab128249; 1:500) and rabbit anti-phospho-JNK1/JNK2 (Thr183, Tyr185) (ThermoFisher MA5-14943; 1:500). Secondary antibody was goat anti-mouse AlexaFluor 647 at 1:1000 (ThermoFisher). Images were taken with a Zeiss LSM 980 confocal microscope with an EC ‘Plan-Neofluar’ 20x/0.50 M27 air objective, taking z-stacks through the entire brain with step sizes of 1-2 μm. Images were visualised and analysed using ImageJ. TH+ cells were manually counted. For fluorescence quantification, z-stacks of the whole brain were 3D-projected using a maximum intensity projection. From these whole-brain stacks, ROIs were drawn around the horizontal mushroom body lobes, an axonal/synaptic neuropil region receiving dopaminergic input. Mean fluorescence values were taken with background subtracted. Brains compared were dissected, stained and imaged in parallel.

### Locomotor and sleep analysis

#### DAM

Sleep analysis was performed using the *Drosophila* Activity Monitor 5 system (DAM5H, Trikinetics inc., MA, USA), which improves on the fidelity of the earlier DAM2 system by using 15 infrared beams per tube, rather than a single one, to track movement ^68^. Male flies of the age indicated were loaded into glass behavioural tubes (Trikinetics inc., MA, USA) containing 4% sucrose and 2% agar and left for two full days to acclimatise. On the third day the number of beam breaks was measured using *Drosophila* Activity Monitor. Sleep was defined using the standard definition of <u>>5</u> min of inactivity, which has been shown to correlate with postural changes, increased arousal threshold and altered brain activity patterns ^68^. Sleep bout number and duration were analysed using the open-source software suite phaseR (https://github.com/abhilashlakshman/phaseR/tree/main) ^86^.

#### DART

As above, flies were loaded into behavioural tubes with flies. After 24 h acclimatisation, a single stimulus in the form of a mechanical vibration (a train of five 200 ms pulses separated by 800 ms intervals, set at the DART system’s maximum intensity) was delivered at Zeitgeber Time (ZT) 3. Locomotor activity was recorded using the *Drosophila* ARousal Tracking (DART) system (BFKlabs) ^57^ as previously described ^58^. Videos were taken using a USB webcam (Logitech) and speed (mm / s) was analysed by the DART system from absolute position, and binned in 1 min intervals. Flies were excluded from analysis if they exceeded an average of 1 mm / s speed in the five 1 min bins preceding the stimulus. The mean speed (mm / s) in the 1 min bin following the stimulus was analysed.

#### FLLIT

For FLLIT (Feature Learning-based LImb segmentation and Tracking) measurements, de-winged male flies were loaded into custom-made 2x2 cm arenas without anaesthesia. 500 fps videos consisting of flies walking in a straight line and taking at least three clear strides were taken using a Photron FASTCAM Mini UX50 High Speed Camera with a Sigma 105 mm Macro lens. Gait analysis was performed using the FLLIT system ^63^, as previously described ^58^. Where FLLIT generates data for parameters per stride, the first and last stride were excluded and a mean of the remaining values taken. Values for each leg were pooled, giving six values per fly. Definitions of the parameters used are as follows. *Stride length*: Straight-line distance between the posterior and anterior extreme positions of the limb in each stride. Directly output by FLLIT as “Displacement”. *Path / displacement*: Path Travelled (directly output by FLLIT) is the total actual distance covered by a limb during a stride; this is divided by Stride Displacement to compensate for differences in stride length. Larger values indicate greater deviation from an optimal straight-line path. *Movement %*: % of the video for which a given limb is in motion.

### Quantification and statistical analysis

Statistical analyses were performed using Graphpad Prism. Data sets were tested for normal distributions using the Shapiro-Wilk test. Normally distributed data sets were tested for statistical differences using unpaired t-tests with Welch’s correction for non-identical variance, and non-normally distributed data sets were tested using Mann-Whitney U-test. Interactions between genotype and age/food type were analysed using 2-way ANOVA with Sidak’s correction for multiple comparisons.

## ACKNOWLEDGEMENTS

This study was funded by a MRC Senior Non-Clinical Fellowship (MR/V03118X/1) to J.E.C.J, a BBSRC Project Grant (BB/X00094X/1) to S.L and J.E.C.J.

## AUTHOR CONTRIBUTIONS

S.A.L – Conceptualization, Methodology, Investigation, Writing - Original Draft, Writing – Review & Editing, Visualization, Funding acquisition; J.E.C.J – Writing – Review & Editing, Supervision, Project administration, Funding acquisition.

## DECLARATION OF INTERESTS

The authors declare no competing interests.

## SUPPLEMENTARY FIGURE LEGENDS

**Supplementary Fig. 1.**

Two representative whole ex vivo brains of 20 d post-eclosure control flies raised on food contaminated with 100ppm fluorophore (540/560)-tagged 0.1 μM polystyrene microspheres. Arrows indicate accumulations of MNPs. Scale bar = 50 μM. Note: left image is also shown in Fig. 1A.

**Supplementary Fig. 2.**

**A-B.** Manually counted TH^+^ cell bodies within the PPLM1+2 (**A**) and PPM3 (**B**) subpopulations. Grey bars indicate control and blue bars *park*^1^ flies. Empty bars indicate flies were raised and maintained on std food; filled bars indicate MNP-containing food. Comparisons are 2-way ANOVA, within genotype, with age (young = 3-5 d; aged = 30 d) and diet (std vs MNP-containing food) as factors. *P < 0.05.

**Supplementary Fig. 3.**

**A.** Schematic showing the use of the *Drosophila* ARousal Tracking (DART) system to assay stimulus-dependent locomotion.

**B.** Traces showing average speed (mm / s) over time in 1 min bins for control and *park*^1^ 10 d old flies, in standard vs MNP food, in response to a mechanical stimulus occurring at 10 mins.

**C.** Age-dependent change in motor performance. Quantification of mean speed in the 1 min bin immediately following mechanical stimulus, at 4 age points. *N* values indicated on graph, referring to number of animals. Comparisons are 2-way ANOVA, within genotype, with age (3, 10, 20 and 30d post-eclosure) and diet (std vs MNP-containing food) as factors. **P < 0.01.

**Supplementary Fig. 4.**

Quantification of the FLLIT-derived gait parameter movement %. Data points represent individual limbs, 6 per fly. Comparisons are 2-way ANOVA, within age point, with genotype and diet as factors. *P < 0.05, **P < 0.01, *** P < 0.001, ****P < 0.0001.

## Notes

### Competing Interest Statement

The authors have declared no competing interest.

### Summary of Updates

Updated version; author affiliations have been corrected.

