## Supplementary figures and images for "Micro-/nano-plastics accentuate Parkinson’s Disease-relevant phenotypes in a *Drosophila* model"

### Supp. Fig 1

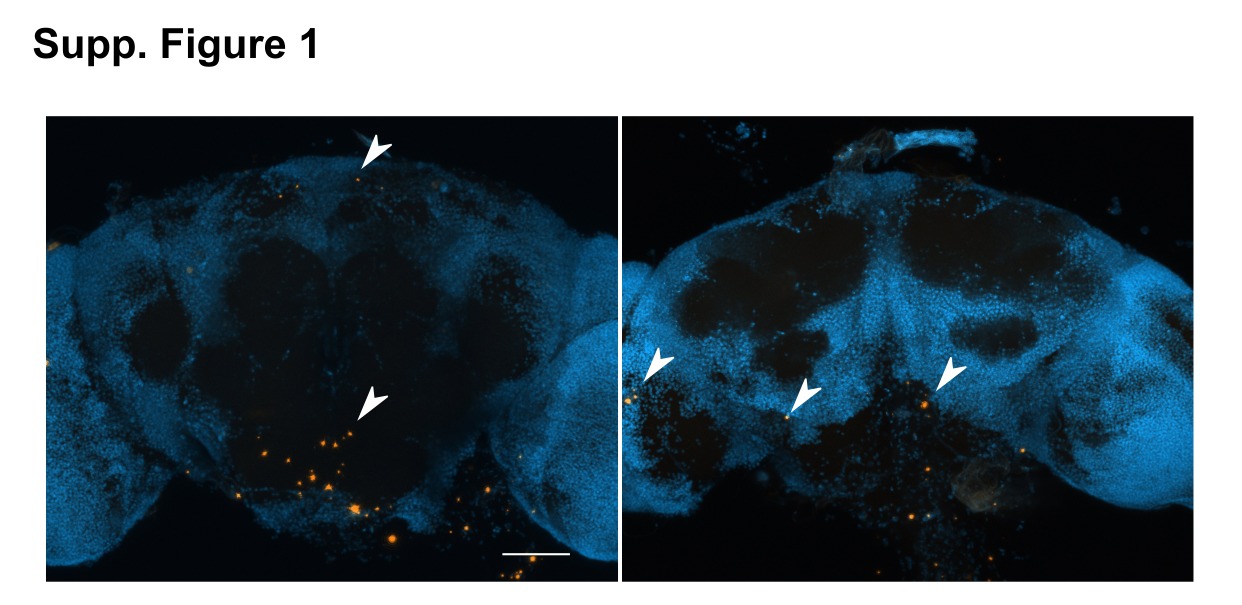

### Supp. Fig 2

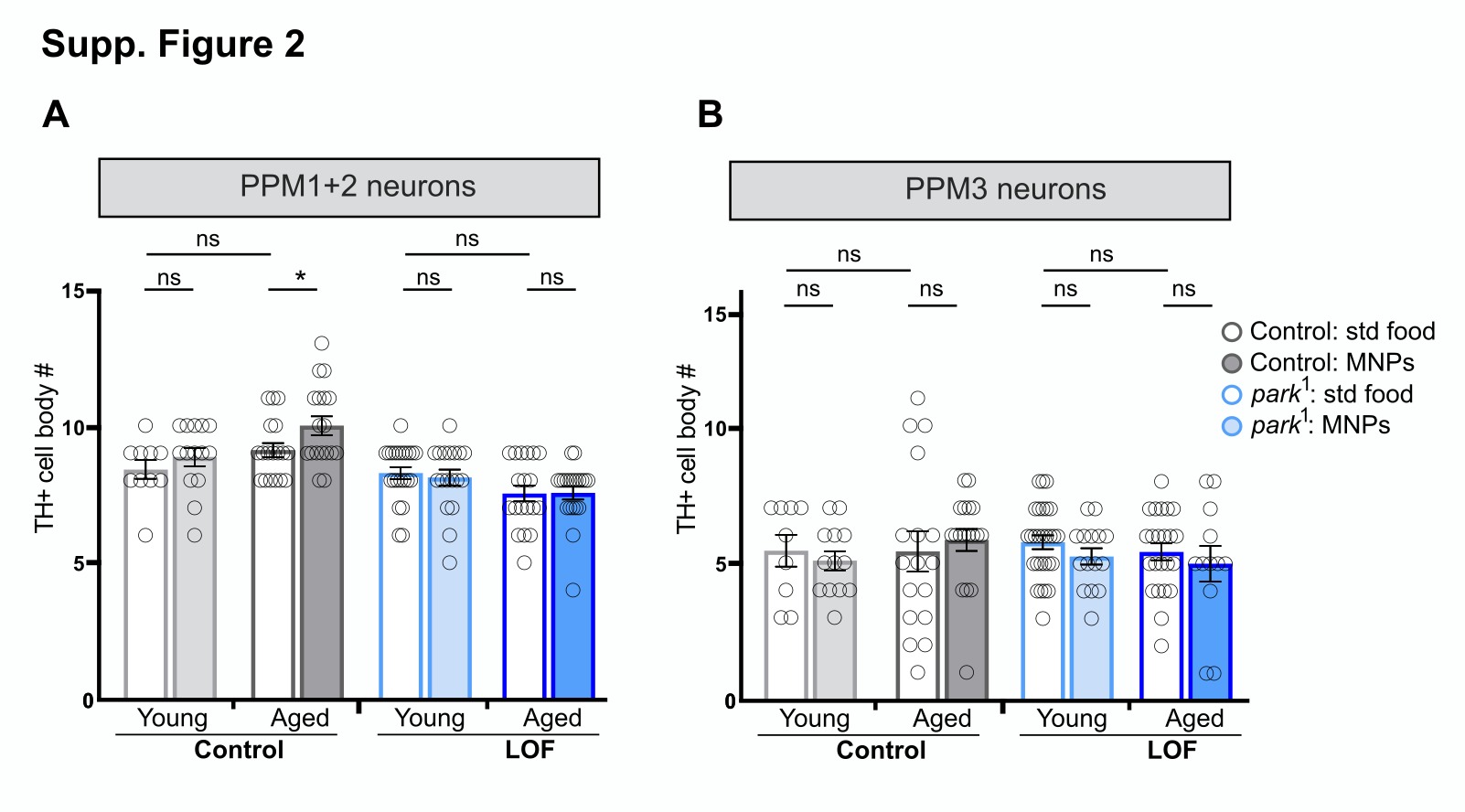

### Supp. Fig 3

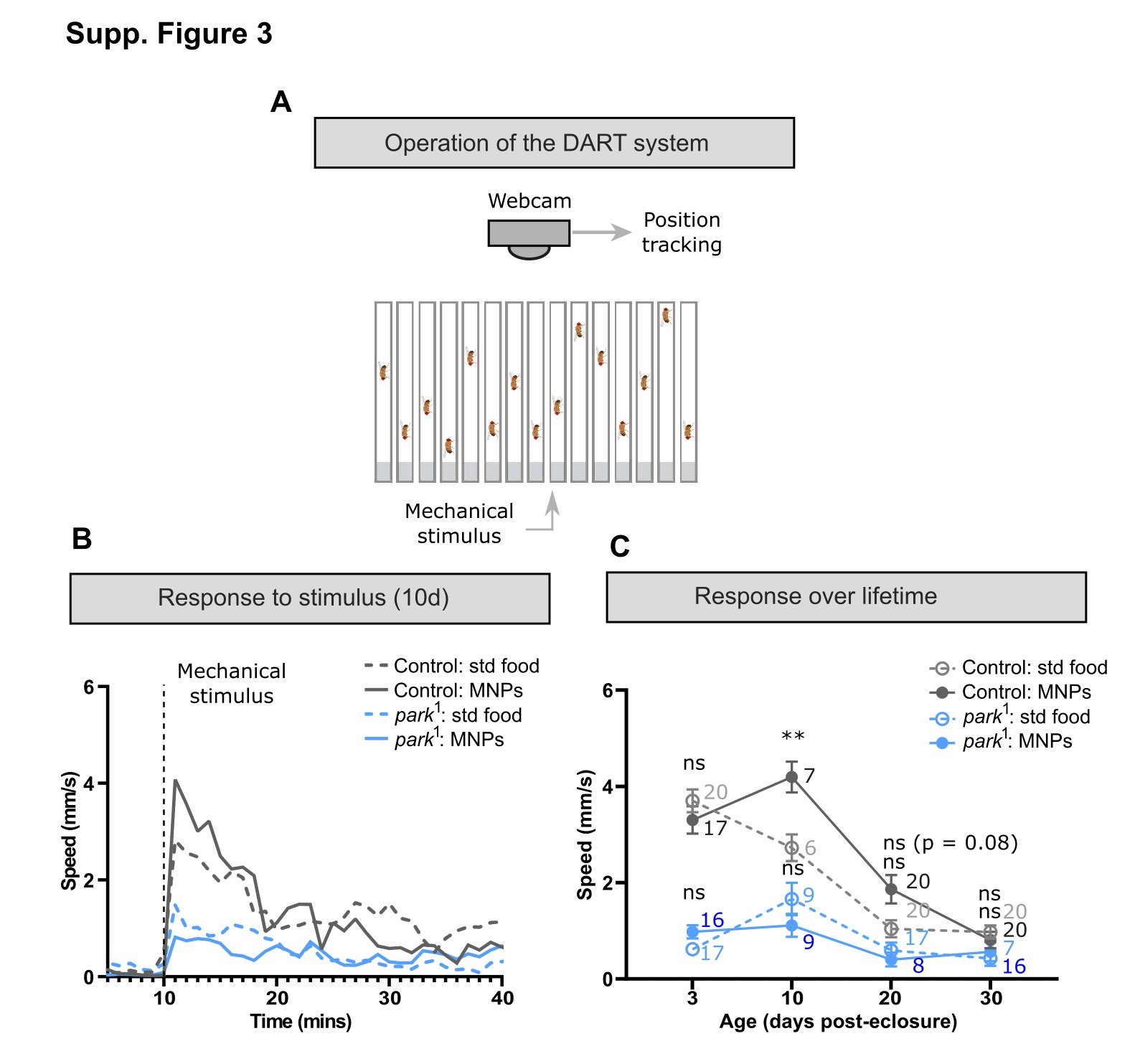

### Supp. Fig 4

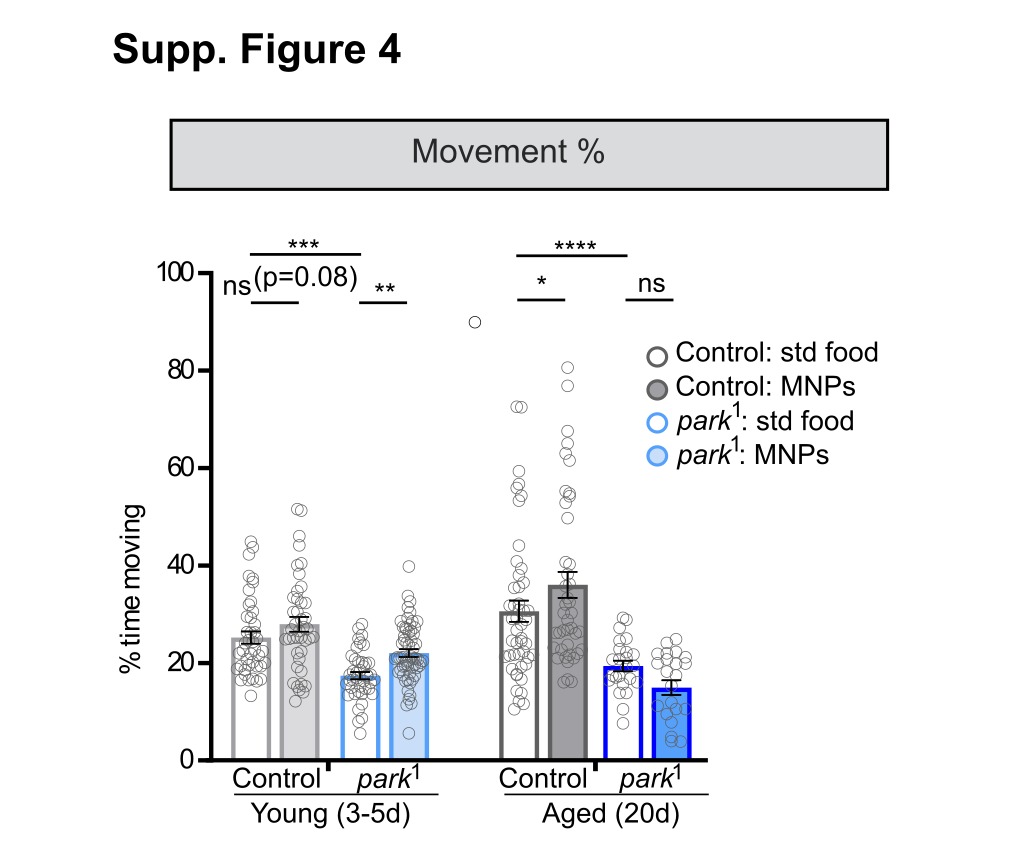
